# Application of a High-Biomimetic Tumor Organoid-CAF Co-Culture Model for the Efficacy Evaluation of CAR-T Drugs

**DOI:** 10.64898/2026.04.16.718819

**Authors:** Jiagen Li, Jian Wang, Yan Sun, Jianchuang Liu, Lijie Rong, Rongrong Xiao, Xiaoni Ai

## Abstract

The tumor microenvironment (TME) is a complex ecosystem composed of tumor cells, cancer-associated fibroblasts (CAFs), immune suppressive cells, and the extracellular matrix (ECM), playing a crucial role in tumor development and CAR-T cell therapy efficacy. CAR-T therapy has shown promise in hematological malignancies but faces challenges in solid tumors due to the TME’s ability to suppress CAR-T cell infiltration, proliferation, and cytotoxicity. Traditional drug evaluation models, such as 2D cell cultures and animal models, have significant limitations due to oversimplification of the in vivo environment or physiological differences between species. Organoid models offer a more biomimetic approach but often fail to fully recapitulate the TME’s complexity and heterogeneity. Our research developed a tumor organoid and CAF co-culture model using the IBAC co-culture chip, demonstrating that CAFs significantly impact CAR-T cell therapy efficacy by forming physical (e.g., fibronectin) and chemical (e.g., IL-10) barriers that prevent CAR-T cell infiltration and cytotoxicity. This model provides a high-biomimetic platform for investigating the TME’s effects on CAR-T therapy and highlights the importance of incorporating a comprehensive stromal component into in vitro models to enhance their predictive power for cancer treatment.

## 1. Introduction

The tumor microenvironment (TME) is a complex micro-ecosystem composed of tumor cells and surrounding non-tumor cells, including cancer-associated fibroblasts (CAFs), immune suppressive cells, and the extracellular matrix (ECM).^[1]^ The TME plays a crucial role in tumor development, progression, and treatment response. Chimeric antigen receptor (CAR)-T cell therapy is a novel immunotherapy approach that involves genetically modifying a patient’s T cells to express a CAR, allowing them to recognize and eliminate tumor cells.^[2]^ While CAR-T cell therapy has shown significant therapeutic efficacy in treating hematological malignancies, it still faces significant challenges in treating solid tumors.^[3]^ Current research suggests that the TME can suppress CAR-T cell infiltration, proliferation, and cytotoxicity, thereby reducing the therapeutic effectiveness of CAR-T cell therapy.^[4]^ CAFs are the most prominent cellular component of the TME, and they can secrete large amounts of cytokines and ECM, forming physical, biological, and chemical barriers that prevent CAR-T cell infiltration and cytotoxicity.^[5]^ The ECM, as an essential component of the TME, can also reduce its cytotoxicity by physically obstructing CAR-T cell movement.^[6]^ Therefore, it is essential to consider the influence of the TME when evaluating the efficacy of CAR-T cell therapy.

Traditional drug efficacy evaluation models, such as 2D cell cultures and animal models, suffer from significant limitations. 2D cell cultures oversimplify the in vivo environment by lacking the complex 3D architecture, cell-cell interactions, and heterogeneity found in tissues, leading to poor prediction of clinical outcomes. Animal models, while providing an in vivo context, are hampered by species differences in physiology, metabolism, and immune response, raising ethical concerns, incurring high costs, and exhibiting limited translatability to human patients, especially for complex diseases.

Current in vitro models for evaluating CAR-T cell therapy often lack and neglect the role of CAFs, leading to biased results that do not reflect the actual situation.^[7]^ The lack of a diverse and functional stroma, including fibroblasts secreting extracellular matrix, impacts the physical and biochemical cues that influence tumor cell behavior, drug response, and metastatic potential. The development of highly biomimetic, multicellular models has become an urgent priority. Recently, organoid models have emerged as a promising option. A significant limitation of many current organoid models lies in their incomplete representation of the tumor microenvironment, particularly the absence or underrepresentation of crucial stromal components. Tumor organoid chips are a novel type of in vitro tumor model that can simulate the TME, including tumor cells, CAFs, immune suppressive cells, and ECM. Tumor organoid chips can be used to study tumor development, progression, and treatment response, as well as evaluate the efficacy of anti-tumor drugs.^[8]^ As an emerging research tool, tumor organoid chips have gained increasing attention in recent years for their applications in studying the TME. These chips can mimic the complexity of tumor tissue, including the diversity of cell types and the dynamic process of cell-cell interactions.^[9]^ While some organoid protocols incorporate specific cell types like fibroblasts, the complexity and heterogeneity of the TME are often not fully recapitulated because these materials are typically derived from allogeneic sources and therefore lack patient-specific matching. Without the appropriate stromal interactions, organoids may fail to accurately model processes like immune cell infiltration and efficacy, and matrix remodeling. Furthermore, the absence of key signaling pathways mediated by stromal-derived factors can lead to inaccurate predictions of drug efficacy, as stromal cells can both promote and inhibit tumor growth and therapeutic response. Therefore, incorporating a more comprehensive and dynamic stromal component into organoid models is crucial for enhancing their physiological relevance and predictive power in cell adoptive therapy development. Moreover, current research also highlights some limitations of tumor organoid chips, such as high operational complexity, single evaluative indicators, and insufficient throughput.

The IBAC co-culture chip is a state-of-the-art microfluidic chip that simulates the tumor microenvironment (TME), comprising tumor cells, cancer-associated fibroblasts (CAFs), immune suppressive cells, and the extracellular matrix (ECM). The IBAC co-culture chip offers several advantages: (1) High-throughput: The chip features a 96 micro-well design, enabling the simultaneous cultivation and detection of multiple cell types and test parameters on a single chip. (2) Operational convenience: The open-design of well, and the well diameter are optimized for easy cell seeding and media exchange. This reduces the complexity and labor intensity of manual operation, which in turn enhances the success rate and stability of model construction. (3) High-content whole-well imaging: The chip can be seamlessly integrated with high-content imaging systems, enabling panoramic imaging of entire wells. This significantly increases the information content and resolution of cell observation and analysis, ensuring reliable and accurate data.

We have developed a tumor organoid and CAF co-culture model, which can more accurately simulate the TME. Our research reveals that CAFs can form physical and chemical barriers, preventing CAR-T cell infiltration and cytotoxicity. The physical barrier is mainly composed of ECM, such as fibronectin, while the chemical barrier is formed through the release of immunosuppressive cytokines, including interleukin-10 (IL-10). Our findings suggest that CAFs are a crucial factor influencing CAR-T cell therapy, and their impact should be considered when evaluating the efficacy of CAR-T cell therapy. Tumor organoid and CAF co-culture model provides a new platform for investigating the effects of CAFs on CAR-T cell therapy.

## 2. Results

### 2.1 Establishment of Complex and Highly Mimetic Organoid Co-culture Models

Lung cancer is a complex and heterogeneous disease, characterized by a tumor microenvironment composed of multiple cell types, including tumor cells, CAFs, and tumor-infiltrating immune cells.^[10]^ CAFs play a crucial role in tumor growth, invasion, and immune evasion.^[11]^ In this study, we obtained tumor tissue from lung cancer patients generating lung cancer tumor organoids (LCOs) and paired CAFs. And then we established a co-culture model of tumor organoids and CAFs, mimicking the cell-cell interactions in the tumor microenvironment to evaluate the cytotoxic effects of CAR-T cell therapy (**Figure 1**). The depicted workflow begins with the preparation of lung cancer tissue, from which CAFs and organoids are isolated and cultured separately. The CAFs and organoids are then utilized to establish two distinct models: a co-culture model and a mono-culture model. In the co-culture model, CAFs are combined with organoids, while in the mono-culture model, organoids are cultured alone. Subsequently, these models are employed for the assessment of chimeric antigen receptor (CAR)-T cell therapy. CAR-T cells are introduced to the models to evaluate their interaction with the cancer cells within the TME. After the CAR-T assessment phase, the samples undergo high-throughput and flow cytometry readout, which involves analyzing the outcomes of the CAR-T cell therapy on a large scale. Finally, the data collected from the readouts are subjected to intelligent analysis. This involves using advanced analytical tools to interpret the data, with the aim of gaining insights into the efficacy of CAR-T cell therapy and understanding how it interacts with different components of the TME. The intelligent analysis helps in visualizing and quantifying the therapeutic effects over time, providing a comprehensive understanding of the treatment’s potential.

**Figure 1.**
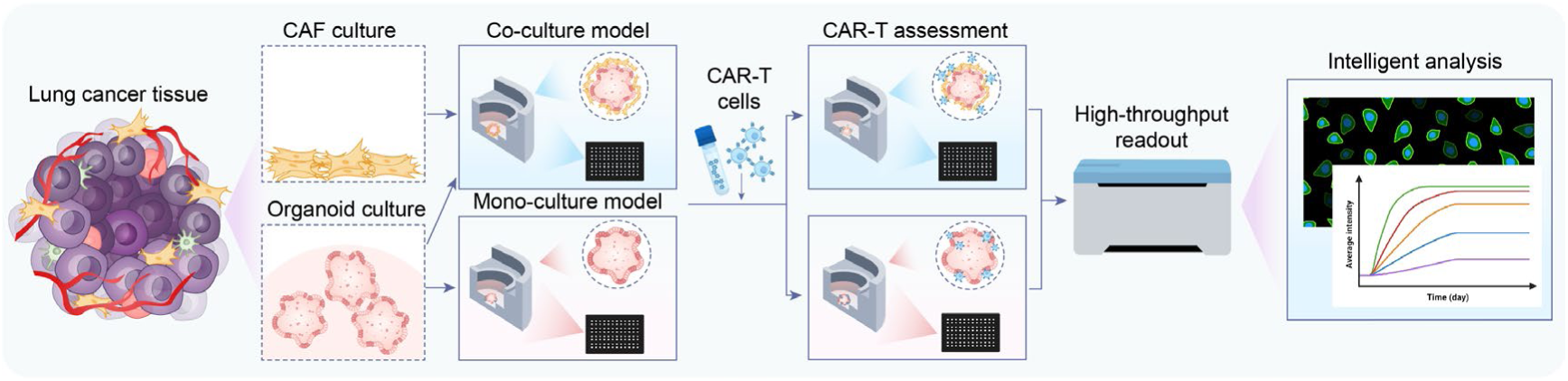
Schematic representation of the experimental workflow for mono-culture and co-culture systems with CAR-T cells. This schematic illustrates the process of culturing tumor organoids and cancer-associated fibroblasts (CAFs) with CAR-T cells. The workflow begins with the resection of tumor tissue, followed by culture of organoids and CAFs respectively. They are then plated for organoids mono-culture model and co-culture models with CAFs. Then, all the models were co-cultured with CAR-T cells. Finally, the level of apoptosis is assessed to evaluate the effectiveness of the CAR-T cells within these platforms.

### 2.2 Histological and Molecular Characteristics of LCOs

In this study, we have successfully extracted fibroblasts from the remaining tissue of the organoids and have successfully cultured them. This process employed precise adhesion selection and conditional culture medium technologies, thereby ensuring the extraction of pure and viable CAF cells from the complex tumor microenvironment. Finally, we enrolled 10 patients, including, 6 matched pairs of organoids and CAFs, along with an additional 4 unmatched CAF samples for further characterization. We first performed a comprehensive analysis of the constructed tumor organoids and their corresponding primary tumor tissues, focusing on morphological and molecular genetic consistency. H&E staining revealed that the tumor-derived organoid samples exhibited patient-specific heterogeneous morphologies. Subsequent pathological marker expression analysis showed that the organoid samples and their corresponding primary tumor tissues shared highly similar characteristics in terms of CK7, TTF-1, and Ki-67 staining patterns (**Figures** 2A and S1A). Immunofluorescence staining results also confirmed these consistent findings (Figure 2B and S1B). Meanwhile, the organoids derived from cancer patients also recapitulated the genomic characteristics of the corresponding tumors, including DNA mutations and copy number variations (CNVs). We performed whole-exome sequencing (WES) on the five organoid samples and their corresponding primary tumors to validate these observations. The total mutation burden and point mutation types were similar between the organoid samples and the paired tumors. Gene expression correlation analysis revealed that the PDO samples retained highly similar gene expression profiles to those of the paired tumors (Figure 2C). Finally, we analyzed the mutation status of commonly mutated genes in lung cancer. The results showed that the mutation status of clinically high-frequency mutated genes, such as TP53, KRAS, and ALK, was highly consistent between the lung cancer organoid samples and primary tumor tissues (Figure 2D). We compared the genome-wide CNVs between the organoid samples and paired tumors, and found that DNA copy number losses and gains were preserved across the genome (Figure 2E and S2C). Overall, these data indicate that the organoid samples recapitulated the histological features and marker expression profiles of the original tumor tissues, consistent with previous reports. These findings collectively demonstrate the accuracy of our organoid sample model in recapitulating the molecular and genetic characteristics of primary lung cancer tissues. Using patient-derived tumor organoids as preclinical models can more accurately reflect disease conditions, promote the development of personalized treatment strategies, and increase the efficiency of drug development.

**Figure 2.**
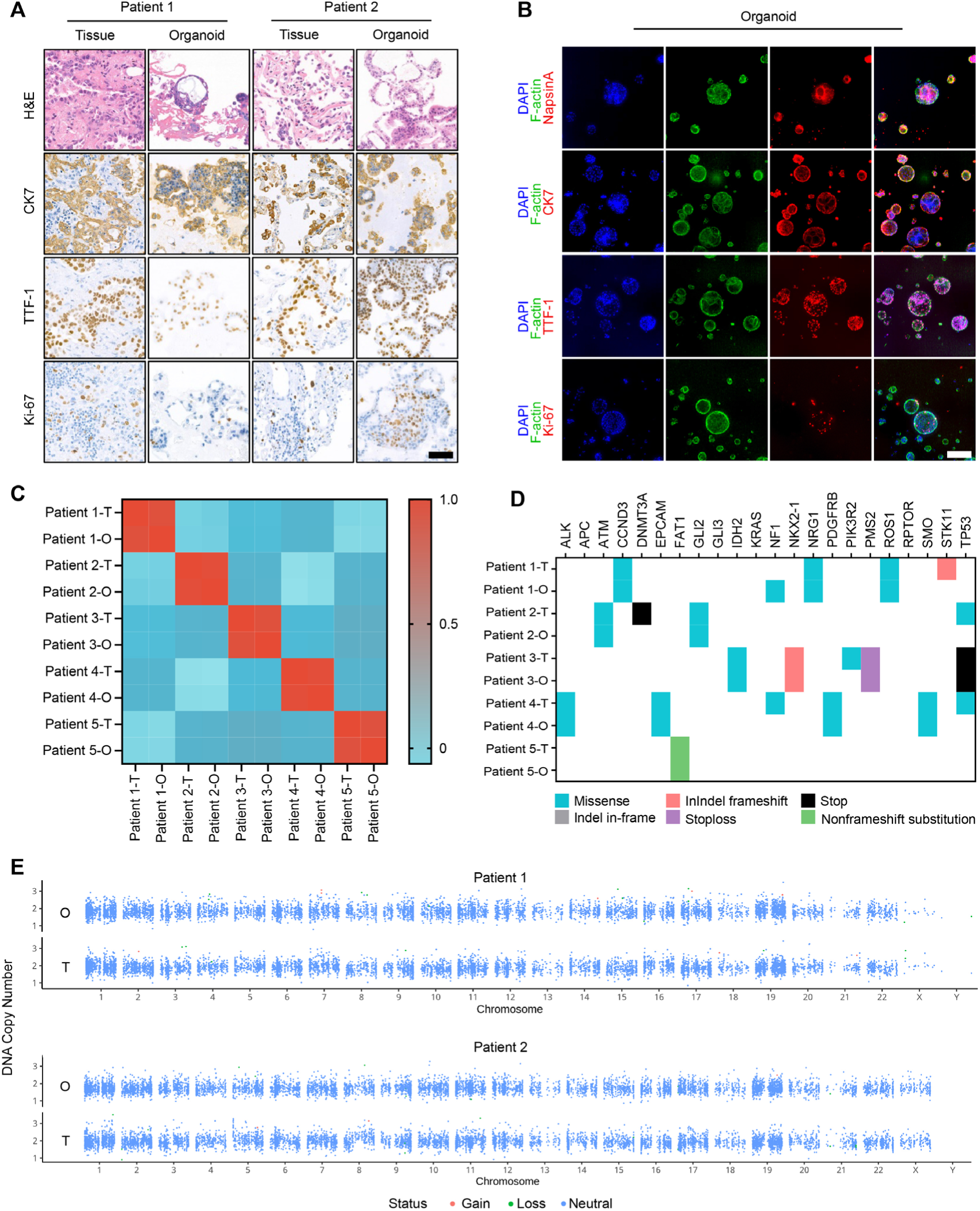
Histological and Molecular Concordance Analysis of Tumor Tissue and Organoid. (A) Histological and immunohistochemical concordance of tumor tissue and organoids from Patient 1 and Patient 2. H&E staining shows the overall histological structure, while IHC for CK7, TTF-1, and Ki-67 demonstrates the expression of these markers in both tissue and organoid samples. Scale bars represent 100 μm. (B) Fluorescence microscopy images of Patient 6 organoids. DAPI staining (blue) highlights the nuclei, F-actin staining (red) shows the cytoskeleton, and immunofluorescence for NapsinA, CK7, TTF-1, and Ki-67 (green) indicates the expression and localization of these proteins. Scale bars represent 100 μm. (C) Heatmap showing the correlation between gene expression profiles of tumor (T) and organoid (O) samples from five patients. The color scale ranges from blue (low correlation) to red (high correlation), indicating the similarity in gene expression between corresponding tumor and organoid samples. (D) Mutation profile comparison between tumor (T) and organoid (O) samples. (E) DNA copy number variation (CNV) analysis of Patient 1 and Patient 2 between tumor (T) and organoid (O) samples.

### 2.3. Functional Characterization of Primary CAFs and Construction of Co-Culture model

Under the microscope, CAF cells, as a type of mesenchymal cell, exhibited unique stellate or spindle-shaped morphologies, which were particularly evident under bright-field capture (Figure S2A). To accurately identify these cells, we detected three specific markers for CAFs. The results showed that FAP (fibroblast activation protein), α-SMA (α-smooth muscle actin), and Vimentin were strongly expressed in CAF cells from different patients (**Figures** 3A and S2B), further confirming that we successfully isolated and cultured CAF cells. In addition to morphological and marker identification, we also investigated the functions of CAF cells in the tumor microenvironment. One important feature of CAF cells is their ability to secrete multiple growth factors, which have significant impacts on tumor growth, invasion, and metastasis.^[1b, 12]^ Therefore, we utilized ELISA technology to detect the growth factors secreted by these CAF cells. The results showed that these CAF cells could secrete various growth factors, including HGF (hepatocyte growth factor), VEGF-A (vascular endothelial growth factor A), and FGF-7 (fibroblast growth factor 7) (Figure 3B). These findings are consistent with previous research reports, further demonstrating that CAF cells can influence tumor occurrence and development through complex factor regulatory effects.^[11c, 13]^ In summary, we successfully constructed the paired lung cancer tumor organoids and CAF cells, and performed comprehensive morphological, molecular genetics, and functional analyses on these cells.

**Figure 3.**
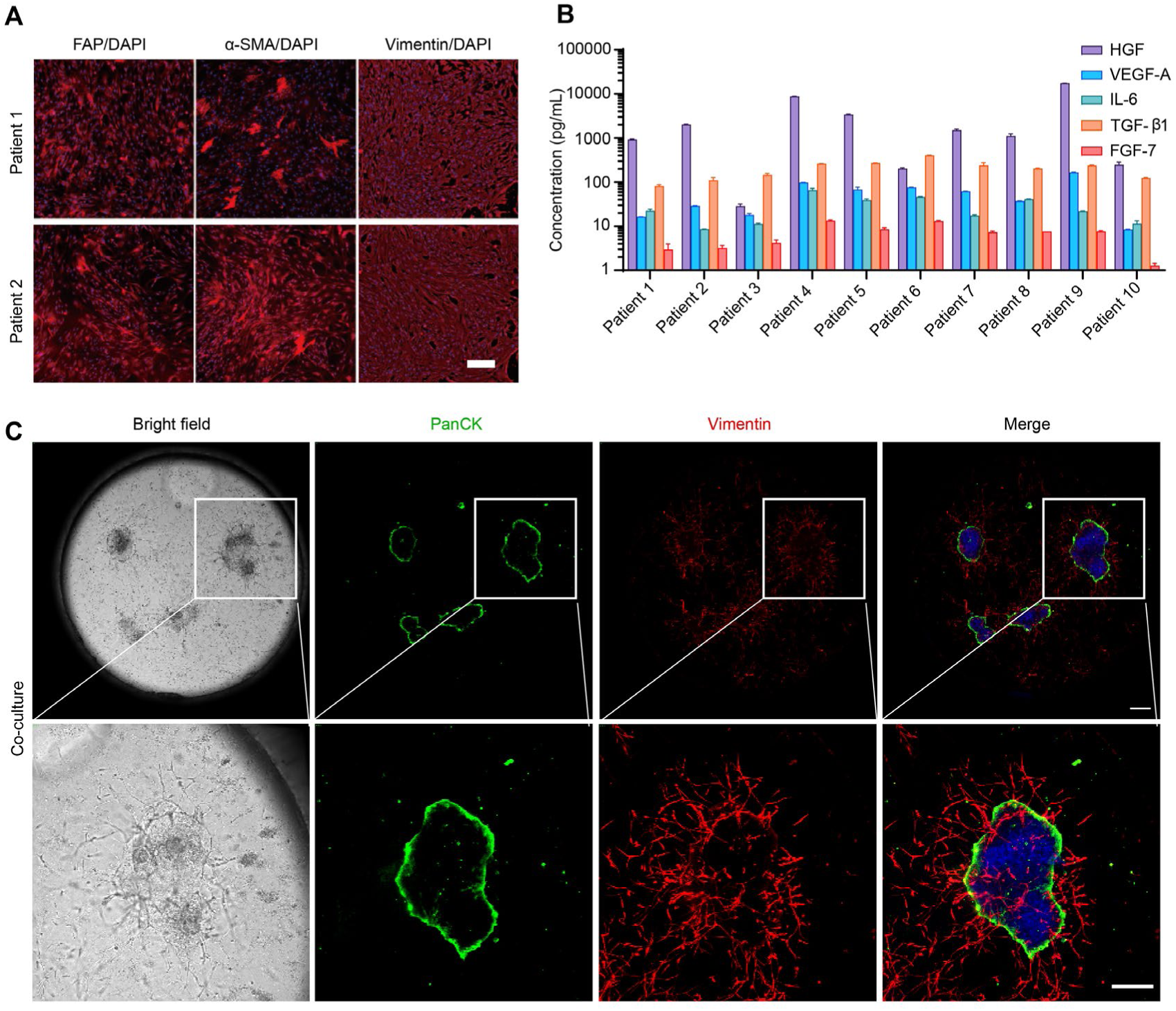
Construction of Organoid Co-Culture Models with CAFs. (A) Immunofluorescence staining of patient-derived CAFs showing the high expression of FAP, α-SMA, and vimentin. Scale bar represents 100 μm. (B) Quantification of cytokine and growth factor concentrations in conditioned media from patient-derived CAFs cultures. Levels of HGF, VEGF-A, IL-6, TGF-β and FGF-7 were measured across different patient samples. Concentrations are presented on a logarithmic scale. (C) Co-culture of organoid and CAF cells. Bright field image shows spheroids, while immunofluorescence staining highlights PanCK (green) and vimentin (red). Scale bar represents 50 μm.

In the latest in vitro studies of CAR-T, investigators often focus on single-culture models of tumor organoids, overlooking the significant influence of other components in the tumor microenvironment.^[14]^ Among these components, CAFs play a crucial role in tumor occurrence and development, accounting for a high percentage approximately 15%-85% within tumor stromal cells.^[11c, 15]^ To more comprehensively simulate the tumor growth microenvironment, we innovatively employed paired lung cancer organoids and CAF cells to construct an LCO+CAF co-culture model. During the construction of this model, we utilized immunofluorescence technology to perform a detailed characterization. From the captured images, we can clearly observe that, under co-culture conditions, organoids and CAF cells express their respective markers. The CAF cells rapidly enveloped the organoids, forming a larger, more irregular 3D structure (Figure 3C). This structure resembles a miniature tumor tissue, exhibiting characteristics more closely resembling those of real tumor growth. The advantages of co-culture model are primarily reflected in two aspects. Firstly, by introducing CAF cells into the model, we successfully simulated the interactions between CAF cells and tumor cells during in vivo tumor growth, thereby increasing the reductionism of tumor model. Secondly, since we used paired lung cancer organoids and CAF cells from the same patient, we can effectively reduce cellular heterogeneity that may arise from constructing models with unmatched tissues. This high level of matching and consistency enables our model to more accurately reflect the tumor characteristics of individual biological entities. In summary, this LCO+CAF co-culture platform provides us with a more comprehensive and bio-mimetic in vitro research.

### 2.4. Evaluating CAR-T Cell Therapy Efficacy Using an LCO+CAF Co-Culture Platform

Clinical trial data show that CAR-T cell therapy achieved a limited success rate, providing challenge for further research and application.^[16]^ Besides, considering the high costs and time-consuming nature of clinical trials, we seek a more efficient and economical method to assess CAR-T product efficacy. In this study, selecting a highly bio-mimetic and in vitro tumor model for preclinical CAR-T product efficacy evaluation becomes crucial. Based on this idea, we constructed a unique co-culture model and used it to evaluate the cytotoxicity of CAR-T cells against paired LCOs and CAFs from two patients. Before the experiment, we performed a detailed characterization of the mesothelin target expression in the two LCOs and CAF lines using IHC technology. We confirmed robust mesothelin expression in both LCO models, with expression levels mirroring those observed in primary tumor specimens, while CAF populations exhibited far lower mesothelin expression (**Figures** 4A and 4B). To verify whether the antigens expressed on the organoids and CAF cells have biological function, we co-cultured the aforementioned organoids and CAF cells with CAR-T cells respectively and assessed the proliferation and activation status of the CAR-T cells using flow cytometry. The results indicated that the organoids indeed promote the activation and proliferation of CAR-T cells, whereas CAF cells exhibited no detectable stimulatory effects (Figure 4C and S3A). This result provided strong assurance for our subsequent experiments. Next, we utilized fluorescence labeling of caspase 3 to distinguish the killed cells in the co-culture platform. Meanwhile, we employed a high-content imaging system to capture and analyze the apoptosis signals produced in the entire well. The long-term tracking results showed that CAR-T cells exhibited strong cytotoxicity when co-cultured with LCOs alone. However, when we introduced CAF cells to construct a triple cell co-culture model, the tumor-killing ability of CAR-T cells was significantly suppressed (Figure 4D, 4E and S3B). To more intuitively present the microscopic details occurring in these two models, we created real-time animations. In the animations, we can see that during the co-culture of LCOs and CAR-T cells, scattered LCOs were numerous small spheroids and were strongly attacked by CAR-T cells. In contrast, during the triple cell co-culture system, CAF cells acted like a “chain”, tightly gathering LCOs into larger tumor assembloid. This structure reduced the contact between LCOs and CAR-T cells, thereby decreasing the cytotoxic effect of CAR-T cells (Video 1). Through this series of experiments and analyses, we conclude that the LCO+CAF model based on the IBAC co-culture chip not only has a more complex and bio-mimetic structure but also exhibits operational convenience and standardized data analysis advantages, particularly in evaluating CAR-T cell efficacy. The successful construction and application of this model provide a powerful tool for us to further explore the mechanisms of CAR-T cell therapy and offer a reliable toolkit for future preclinical research.

**Figure 4.**
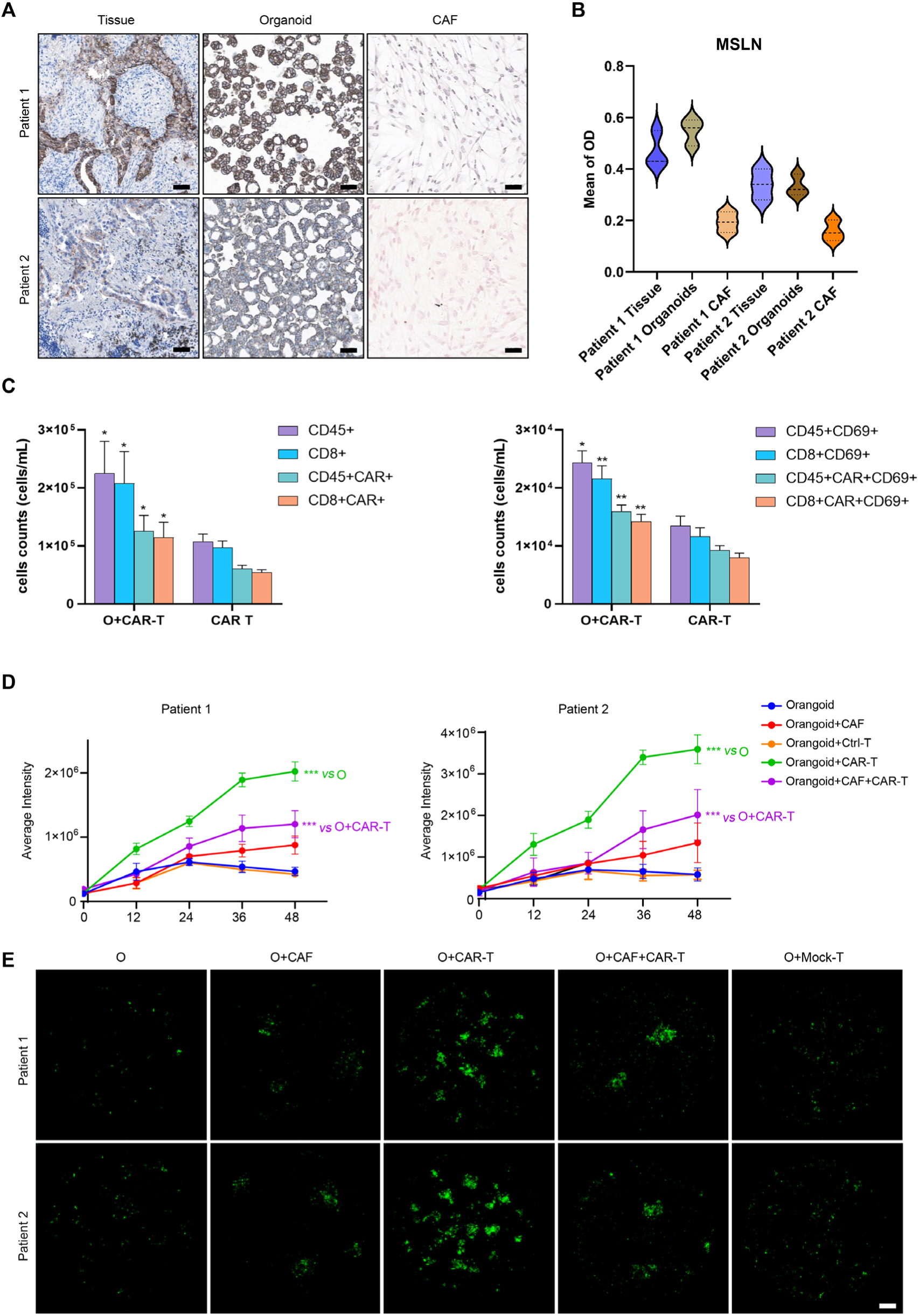
Efficacy evaluation of CAR-T using co-culture model. (A) Mesothelin (MSLN) IHC staining of patient-derived samples including CAF, organoid, and tissue sections. Scale bars represent 100 μm. (B) Violin plots showing the distribution of MSLN expression levels from the same sample types in (A). (C) Bar graphs illustrating the quantification of immune cell populations in various cultivation groups from patient 6. The left graph shows cell counts for different immune cell subsets. The right graph further shows cell counts for activated immune cell subsets. Asterisks indicate significant differences compared to the CAR-T group. (D) Line graphs representing the average intensity of organoid apoptosis over time (0-48 hours) under different cultivation conditions. (E) Caspase-3 fluorescent microscopy images (green) of organoids from (D) at 36h time point. Scale bars represent 100 μm. Significant differences between groups are indicated by asterisks (* = p < 0.05, ** = p < 0.01, *** = p < 0.001).

### 2.5. CAF-Mediated Barrier Function against CAR-T Cell-Mediated Cytotoxicity

CAF cells and T cells were labeled with live-cell fluorescence markers for real-time monitor the interactions between CAR-T cells, LCOs, and CAFs. To further investigate the mechanisms by which the tumor assembloid formation impedes CAR-T cell-mediated cytotoxicity, we employed a series of molecular and cellular characterization tools to systematically analyze the complex. The results revealed that CAF cells exhibited a unique arrangement, encircling LCOs and tightly enveloping them within the complex, which results in reduced CAR-T cell migration (**Figures** 5A and 5B). Besides, we utilized 3D reconstruction image analysis, which demonstrated that CAR-T cell infiltration ability was significantly impaired in the presence of the LCO+CAF tumor assembloid (Figure 5C and 5D). This result further supported the notion that CAF cells exert a barrier function within the complex. To investigate how CAFs establish this biological barrier, we hypothesized that the ECM composition was altered in the co-culture system. Due to primary antibody species limitations, we used double immunofluorescence staining to test this hypothesis, employing vimentin in one channel to label CAFs and the other channel for target protein detection using species-compatible secondary antibodies. Immunofluorescence staining revealed a significantly elevated expression of fibronectin in the co-culture group compared to the mono-culture group (Figure 5E). This observation may partially explain the formation mechanism of the barrier function. In addition to their structural role, we also investigated the biological functional impact of CAF cells. Previous studies have shown that CAF cells release a range of immunosuppressive factors, such as IL-10, which can inhibit the function of effector cells. Therefore, we used quantitative ELISA technology to detect the levels of IL-10 in the co-culture system. The results showed that the secretion of these two factors was significantly increased in the LCO+CAF group compared to the control group (Figure 5F). This suggests that CAF cells further suppressed CAR-T cell-mediated cytotoxicity by releasing immunosuppressive factors. In summary, the LCO+CAF complex suppressed CAR-T cell-mediated cytotoxicity through both the structural barrier function of CAF cells and the release of immunosuppressive factors. These findings provide new research tools for understanding the interactions between cells in the tumor microenvironment.

**Figure 5.**
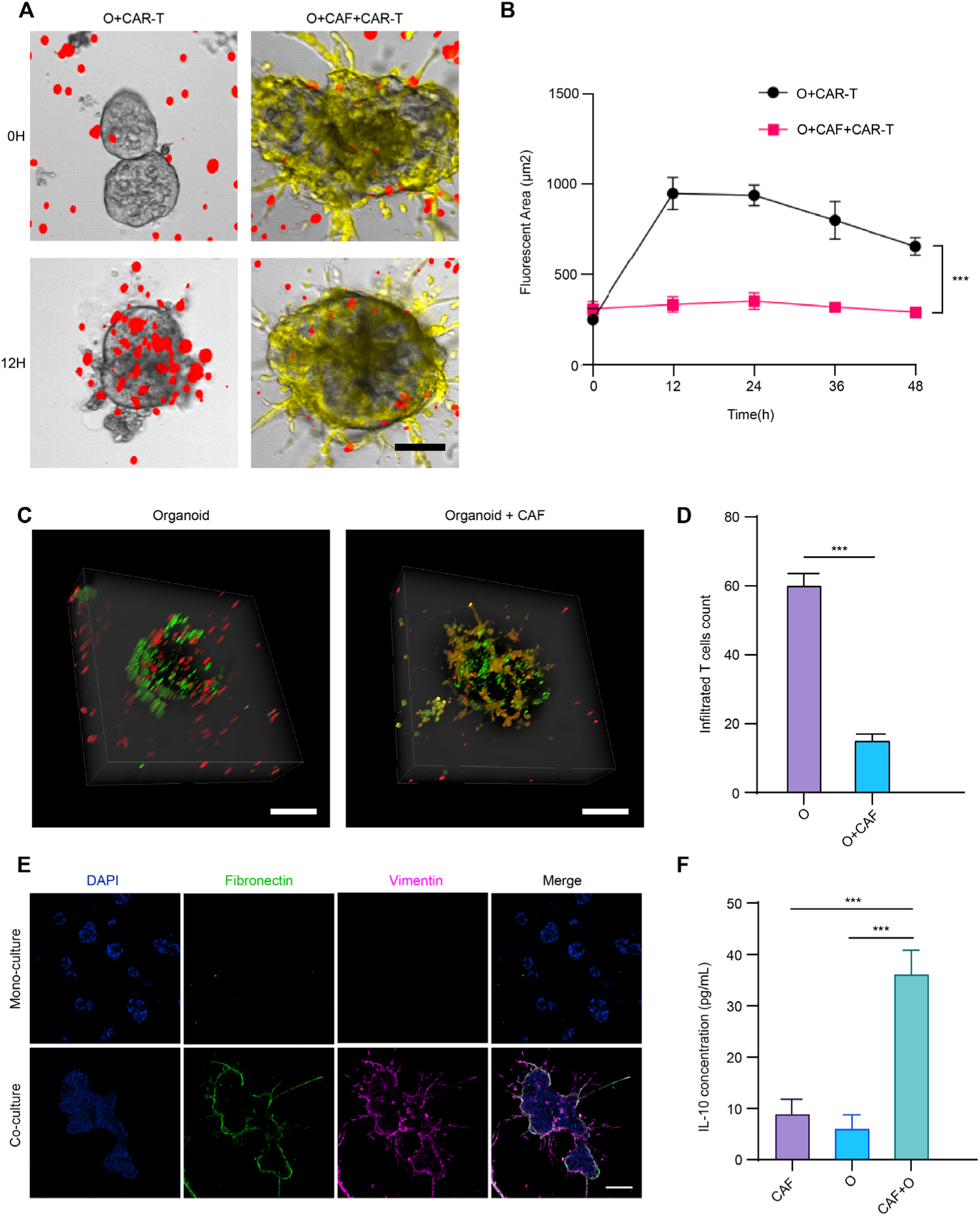
Infiltration and interaction dynamics of CAR-T cells in mono-culture and co-culture models. (A) Confocal microscopy images showing the infiltration of CAR-T cells (red) into organoids (bright field) at 0 hour and 12 hours in the two models. CAF cells are stained yellow. Scale bar represents 50 μm. (B) Quantification of infiltrated CAR-T cells fluorescent area from (A). The graph shows the mean fluorescent area (μm²) over time (0-48 hours) for each condition. (C) 3D confocal microscopy images illustrating the infiltration of CAR-T cells (red) into organoids in the two models. Organoids are shown in bright field, and CAFs are stained yellow. Scale bar represents 50 μm. (D) Quantification of infiltrated T cells from (C). The bar graph shows the mean number of infiltrated CAR-T cells per field in mono-culture and co-culture models. Data are presented as mean ± SD. (E) Immunofluorescence staining showing the expression of fibronectin (green) within co-culture platform. Images are shown for mono-culture and co-culture models. Scale bar represents 50 μm. (F) Quantification of IL-10 concentration in supernatant of different models. Data are presented as mean ± SD. Significant differences between groups are indicated by asterisks (* = p < 0.05, ** = p < 0.01, *** = p < 0.001).

## 3. Discussion

The development of chip-based cell culture and detection platforms has increasingly gained attention and had a profound impact on biomedical research and applications.^[17]^ Our research has demonstrated the potential of chip-based co-culture systems in cancer research, featuring paired CAFs and organoids derived from lung cancer patients. Compared to traditional cell culture methods, this system offers numerous advantages, including high-throughput capability, operational convenience, and high biomimicry. By using this co-culture model, we systematically evaluated the cytotoxicity of CAR-T cells against lung cancer. The results provide important references for assessing the efficacy of CAR-T cell therapy in clinical settings and lay the foundation for further exploration of the tumor microenvironment and its impact on treatment responses.

Traditional organoid co-culture models often use cells from different individuals, which may have inherent limitations, including cellular heterogeneity, immunological issues, low physiological relevance, increased complexity, and difficulty in results analysis.^[18]^ These limitations may compromise the reliability and repeatability of the model. Therefore, it is crucial to select suitable cell sources and control relevant factors to obtain a more physiologically relevant and repeatable in vitro co-culture model.^[19]^ In this study, we used paired LCO and CAF as model construction materials, which can effectively avoid these issues. However, due to the lack of paired immune cells, we performed functional testing using allogenic CAR-T cells.

Our co-culture system utilizing patient-derived organoids and CAFs enables the development of personalized cancer models. By generating co-culture models based on different patients, we can assess the efficacy of various treatment strategies in a personalized setting. This approach has the potential to improve treatment outcomes by determining the most effective treatment method for each patient. Furthermore, the high-throughput capability of our platform enables rapid evaluation of multiple treatment strategies, accelerating the discovery of effective treatment options.

The chip-based co-culture system has a broad impact on cancer research and treatment strategy development. Future studies can utilize this platform to evaluate the efficacy of different treatment strategies, including immunotherapy, targeted therapy, and combination therapy.^[20]^ Moreover, this system can be used to investigate the mechanisms of tumor progression and resistance to treatment, providing valuable insights for developing new treatment strategies. The potential applications of this platform are not limited to cancer research, but also extend to regenerative medicine, tissue engineering, and infectious disease control. Through continuous improvement and optimization of the chip-based co-culture system, we can expect more in-depth research on the tumor microenvironment and personalized medicine, as well as the development of more accurate and effective treatment strategies.

## 4. Experimental Section

### Primary Organoid Establishment and Culture

Fresh lung cancer tumor tissues were washed five times with phosphate-buffered saline (PBS) to remove debris and residual tissue fragments. Fat and blood residues were excised using sterile surgical scissors, and tissues were divided into three portions: one stored at -80°C for whole-exome sequencing (WES), another fixed in 4% paraformaldehyde (P0099, Beyotime) for histopathology and immunohistochemistry (IHC), and the remaining tissue minced into 1–2 mm³ fragments. Tissue fragments were enzymatically digested in Tissue Enzyme Solution Ⅱ (KS100130, Daxiang Biotech) using Intelligent Tissue Dissociator (D8, Daxiang Biotech) under 37°C for 10–15 min. After digestion, mechanical dissociation was performed using 5 mL Organoid Wash Solution (KC100145, Daxiang Biotech) with repeated pipetting (5–10 times). The mixture was filtered through a 100-μm cell strainer (352360, BD Biosciences), centrifuged, and resuspended in lung cancer organoid culture medium (OC100132, Daxiang Biotech). Cells were mixed with 2× volume Matrigel (356231, Corning) and plated into 24-well plates. After polymerization (37°C, 10 min), 500 μL of medium supplemented with anti-apoptotic factor (IA100101, Daxiang Biotech) was added and refreshed every 3–4 days. Organoids were passaged (1:2–1:5 ratio) using Organoid Digestion Solution (100145, Daxiang Biotech) for 3–8 min and reseeded in fresh Matrigel and medium.

The organoid medium is formulated based on the advanced Dulbecco’s Modified Eagle Medium/F12 (DMEM/F12) medium. It is supplemented with a sophisticated blend of active ingredients, which includes: 1X B27, Glutamax, 10 mM HEPES (15630080, Gibco), 100 μg/mL primocin (ant-pm-2, Invivo-Gen), 50 ng/mL recombinant human EGF (AF-100-15, Peprotech), 10 nM gastrin (G9020, Sigma), 500 nM A83-01 (2939, Tocris Bioscience), 1.25 mM N - acetylcysteine (A9165, Sigma), 10 mM nicotinamide (N0636, Sigma), 100 ng/mL recombinant human Noggin (120-10C, Peprotech), and 20% R - spondin1 conditioned media. Additionally, 10 μM Y - 27632 dihydrochloride, a Rho - kinase inhibitor (M1817, Abmole), is incorporated into the culture medium for the initial 2 - 3 days. This carefully composed medium provides the necessary nutrients and regulatory factors to support the growth and maintenance of organoids, with each component playing a crucial role in creating an optimal environment for cellular development and function.

### CAF cells Isolation and Culture

After the tissue was enzymatically hydrolyzed with collagenase, the precipitate was collected through centrifugation or filtration to obtain the desired components. Then, the tissue pieces were evenly coated onto the bottom of the T25 cell culture flask to ensure good contact with the culture medium and facilitate cell adhesion and growth. The flask was then incubated in the incubator for 15-30 min, which allowed the cells to begin to attach and stabilize. Subsequently, 2.5 mL of CAF medium (FC100101, Daxiang Biotech) was slowly added to the flask to avoid sudden changes in the cell environment that might affect cell viability. The culture medium was changed every three days to provide fresh nutrients and remove metabolic waste, maintaining a suitable environment for cell growth. When the CAF cells fusion reaches about 80%, indicating that the cells have grown to a certain density and need more space and nutrients for further proliferation, passaged culture was performed. To avoid contamination of CAF cells, which could affect the purity and experimental results of the cells, cells should be passaged for at least 1 generation before freezing. This allows for a preliminary screening and removal of potential contaminants. Cells in the 2nd-10th generation were used for experimental studies.

### Establishment of the Co-Culture Systems

IBAC co-culture chips were pre-incubated in advance. On the experiment day, Matrigel was added to the co-culture wells at 8 μL per well. The plate was placed in 37°C incubation for 30 minutes to solidify the matrix. Organoids with different growth rates were collected and resuspended in culture medium and seeded into the chip at 1500-3000 cells per well and stable culture for 2-4 days. When constructing the co-culture model of organoids and cancer-associated fibroblasts (CAFs), CAFs were inoculated and labeled with Cell Tracker Orange (C34551, Invitrogen) is done selectively according to the experimental requirements. The detailed description of the relevant operations is as follows:

CAF cells were digested, centrifuged, and resuspended in medium for counting. The required cell quantity was transferred and stained by 37°C incubation for 40 minutes with periodic tube agitation. CAF cells were resuspended in a co-culture medium (1:1 mixed CAF/organoid) and seeded into the chip at 3000-6000 cells/per well. Organoid and CAF status was monitored daily during co-culture, with 300 μL co-culture medium supplementation on the second day. When constructing the triple cell co-culture model of organoids and cancer-associated fibroblasts (CAFs), and T cells (including CAR-T and mock-T), T cells were resuspended in medium containing Cell Tracker Deep Red (C34565, Invitrogen) for staining. T cells added at 1500-3000 cells/per well followed by 5-day continued culture and detection.

A 96-well plate-based co-culture system was established to evaluate CAR-T cell activation in tumor microenvironment-mimicking conditions. Organoids or CAFs from two lung cancer patients were enzymatically dissociated into single cells, seeded at 2.5×10³ cells/well in plates (100 μL/well of organoid or CAF-specific medium). CAR-T cells (thawed 48 hr prior) were added at 5×10⁴ cells/well (200 μL total volume) to three experimental groups: organoid+CAR-T, CAF+CAR-T, and CAR-T alone (control). After 24 hr co-culture, CAR-T cells were collected from supernatants and analyzed by flow cytometry.

### Organoids real-time animations and Apoptosis Quantitation

Lung cancer organoids, with or without CAFs, were co-cultured with CAR-T cells to mimic the tumor immune microenvironment. During the real-time imaging experiment, the co-culture was monitored using a Real-Time Intelligent Organoid Analysis System (B8, Daxiang Biotech) over a period of 6 to 24 hours. Images were captured at 30-minute intervals and subsequently compiled into a time-lapse video. For Apoptosis Quantitation, apoptosis was assessed using the GreenNuc™ Live Cell Caspase-3 Kit (C1168, Beyotime) and completed detection by High-Content Imaging ( ImageXpress Confocal HT.ai, Molecular Devices). Caspase-3 activity was quantified by measuring green fluorescence intensity and normalized to untreated controls.

### Immunohistochemistry (IHC) and quantification

Organoidswere fixed in 4% paraformaldehyde, paraffin-embedded, sectioned, and processed for IHC. Antigen retrieval was performed by boiling in retrieval buffer. Sections were blocked with 3% H₂O₂, respectively incubated with CK7, TTF-1, Ki-67 and mesothelin primary antibodies at 4°C overnight, followed by secondary antibodies and DAB substate (C09-12, OriGene). Nuclei were counterstained with hematoxylin.

Whole-slide imaging of stained tissue sections was conducted using a high-resolution digital scanner. Acquired images were analyzed in HALO™ software (Indica Labs) using the Multiplex IHC module. Regions of interest (ROIs) were manually annotated to isolate specific tissue types. Optical density (OD) thresholds for mesothelin positivity were calibrated using negative controls to differentiate background noise from specific DAB staining. Mean OD was automatically generated for each ROI. Statistical analysis was performed in GraphPad Prism.

### Immunofluorescence (IF)

Organoids and CAF cells were fixed by 4% PFA, permeabilized (0.5% Triton X-100), and blocked with 5% FBS. Primary antibodies of F-actin, panCK, CK7, TTF-1, Ki-67, napsinA, FAP, α-SMA, vimentin and fibronectin were respectively incubated at 4°C overnight, followed by species-specific secondary antibodies (1 h, room temperature). Nuclei were stained with DAPI (ab285390, Abcam). Images were captured using a confocal microscope (Nikon AXR).

### Flow Cytometry

CAR-T cells were harvested and stained with fluorophore-conjugated antibodies (CD45, CD8, CD69, CAR) and matched isotype controls, and analyzed via flow cytometry (BD FACSCelesta). Absolute cell counts were quantified using counting beads (335925, BD). Cells were gated on singlet CD45+CD8+CAR+ populations, and CD69 expression (%) was normalized to assess activation under CAR-T modulated conditions.

*Antibodies*
| Target | Company | Cat No. | Dilution |
| --- | --- | --- | --- |
| CK7 | abcam | ab181598 | 1:100 |
| TTF-1 | abcam | ab227652 | 1:100 |
| Ki-67 | CST | 9449S | 1:1000 |
| Mesothelin | sino biological | 13128-MM12 | 1:100 |
| F-actin<br>(Phalloidin) | Solarbio | SF488 | NA |
| PanCK | CST | 4545T | 1:200 |
| NapsinA | CST | 62434T | 1:200 |
| FAP | CST | 66562S | 1:200 |
| α-SMA | CST | 19245S | 1:200 |
| Vimentin | abcam | ab92547 | 1:200 |
| Fibronectin | abcam | ab2413 | 1:100 |
| Rabbit IgG(H+L) | abcam | ab150080 | 1:1000 |
| Mouse IgG(H+L) | abcam | ab150115 | 1:1000 |
| CD45-BV510 | BD | 563204 | 5 μL/Test |
| CD8-BV421 | BD | 562428 | 5 μL/Test |
| CD69-FITC | Biolegend | 310904 | 5 μL/Test |

### ELISA

Conditioned media from cancer-associated fibroblasts (CAFs) were centrifuged (12,000 ×g, 10 min, 4°C) to remove debris. Supernatants were diluted (1:2–1:10, depending on targets concentration range) and analyzed for HGF, VEGF-A, FGF-7, IL-10, and TGF-β1 using commercial ELISA kits (R&D Systems). Briefly, 100 μL of standards or samples were added to pre-coated wells, incubated (2 h, RT), washed, and incubated with biotinylated detection antibodies (1 h, RT). After washing, streptavidin-HRP (30 min) and TMB substrate (20 min) were added. Reactions were stopped with H₂SO₄ solution, and absorbance (450 nm/570 nm) was measured with microplate reader (Synergy H1, BioTek). Concentrations were calculated against standard curves.

### Whole-Exome Sequencing (WES) and analysis

Organoids from tumor tissues were cultured for 7 days. Both organoids and matched primary tumor tissues had genomic DNA extracted. Paired-end sequencing was done on the Illumina HiSeq platform at Novogene Co., Ltd. (Beijing, China), achieving an average sequencing depth of ≥100 × for tumor samples and matched organoids. Raw reads were quality-checked with FastQC. Adapters and low-quality bases were removed using Trimmomatic. Clean reads were aligned to the human reference genome (GRCh38/hg38) with BWA-MEM. Somatic SNVs and small indels were found using GATK Mutect2. Mutational signatures were analyzed using the Mutational Patterns R package.

## Acknowledgements

J. Li, J. Wang and Y. Sun contributed equally to this work. This study was financially supported by Chinese Academy of Medical Sciences Central Public-interest Scientific Institution Basal Research Fund (2022-RW320-13), the National Key Research and Development Project (2023YFC3505000), Beijing Municipal Science & Technology Commission (Z231100007223001), and the National Natural Science Foundation of China (82422076 and 82174086), Scientific and Technological Innovation Project of China Academy of Chinese Medical Sciences (C12024C002YN), and Key Research and Development Program of Ningxia (2024BEG01006).

## Data Availability Statement

The data that support the findings of this study are available from the corresponding author upon reasonable request.

---

**Figure S1.**
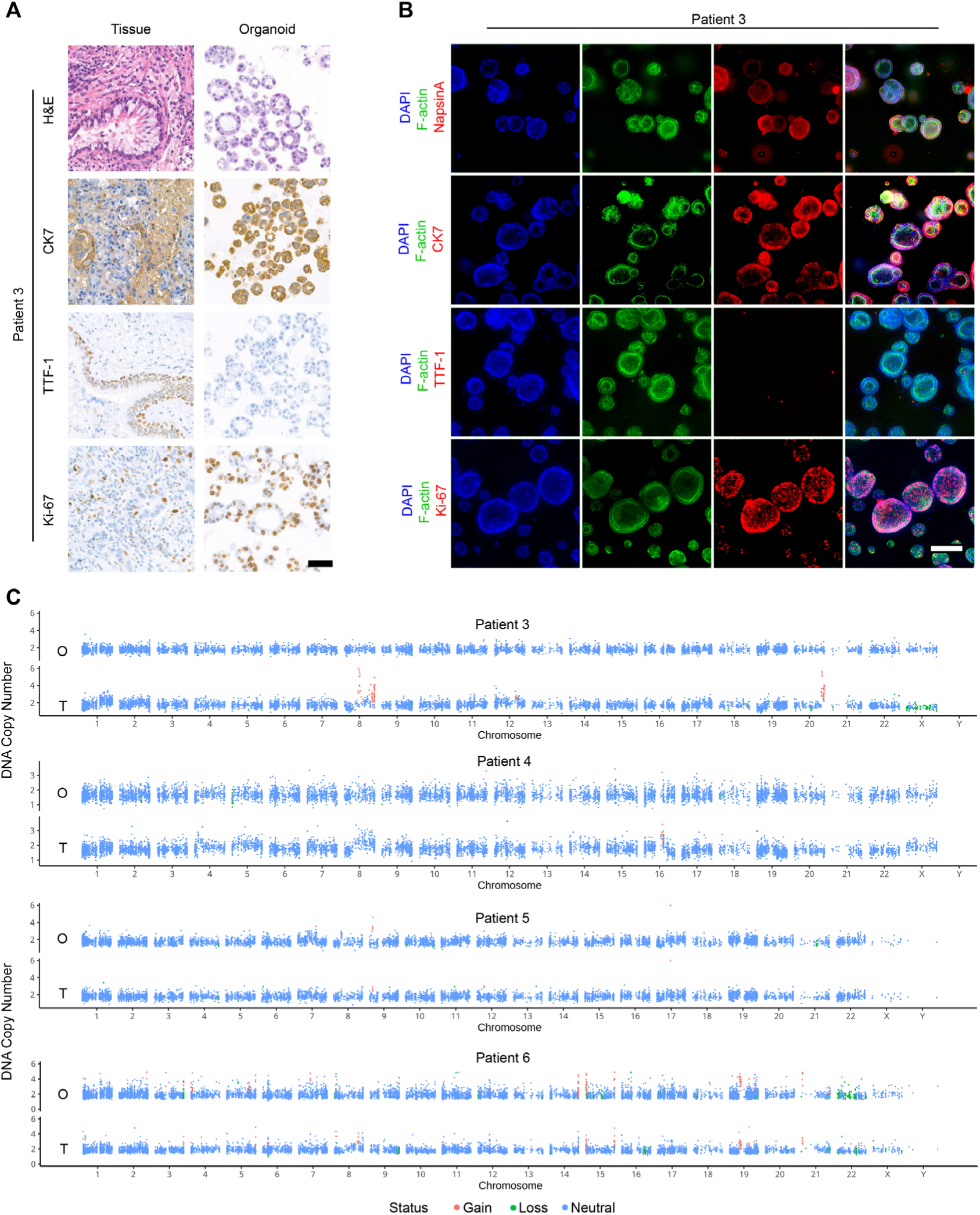
Characterization of patient-derived lung cancer organoids, Related to Figure 2. (A) Histological and immunohistochemical analysis of patient 3 tumor tissue and corresponding organoids. H&E staining shows the overall tissue morphology. Immunohistochemistry was performed to detect CK7, TTF-1, and Ki-67 expression. Scale bars represent 100 µm. (B) Immunofluorescence staining of patient 3 organoids. Organoids were stained for F-actin, Ki67, NapsinA, CK7, and TTF-1. Scale bar represents 100 µm. (C) DNA copy number analysis of four patients-derived organoids using mutation detection. Each point represents a single genomic probe. Red points indicate gains, green points indicate losses, and blue points represent neutral copy number. Chromosomes are indicated along the x-axis.

**Figure S2.**
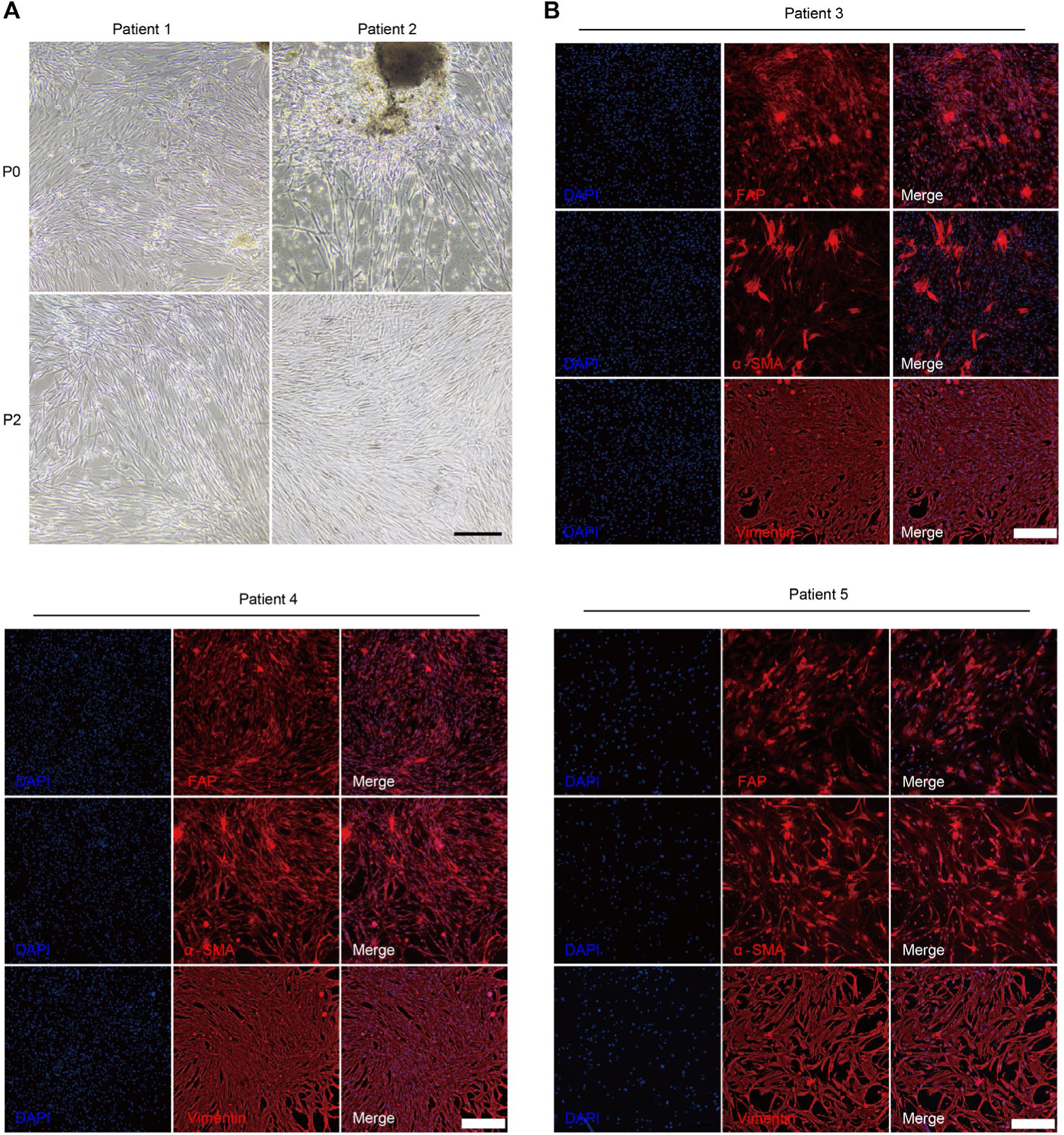
Characterization of patient-derived CAFs, Related to Figure 3. (A) Phase-contrast microscopy images of CAFs from patients 1 and 2 at passage 0 and passage 1. 2. Scale bars represent 200 µm. (B) Immunofluorescence staining of CAFs from patients 3, 4 and 5. Staining shows expression of FAP, α-SMA, and vimentin. DAPI staining indicates nuclei. Scale bar represents 100 µm.

**Figure S3.**
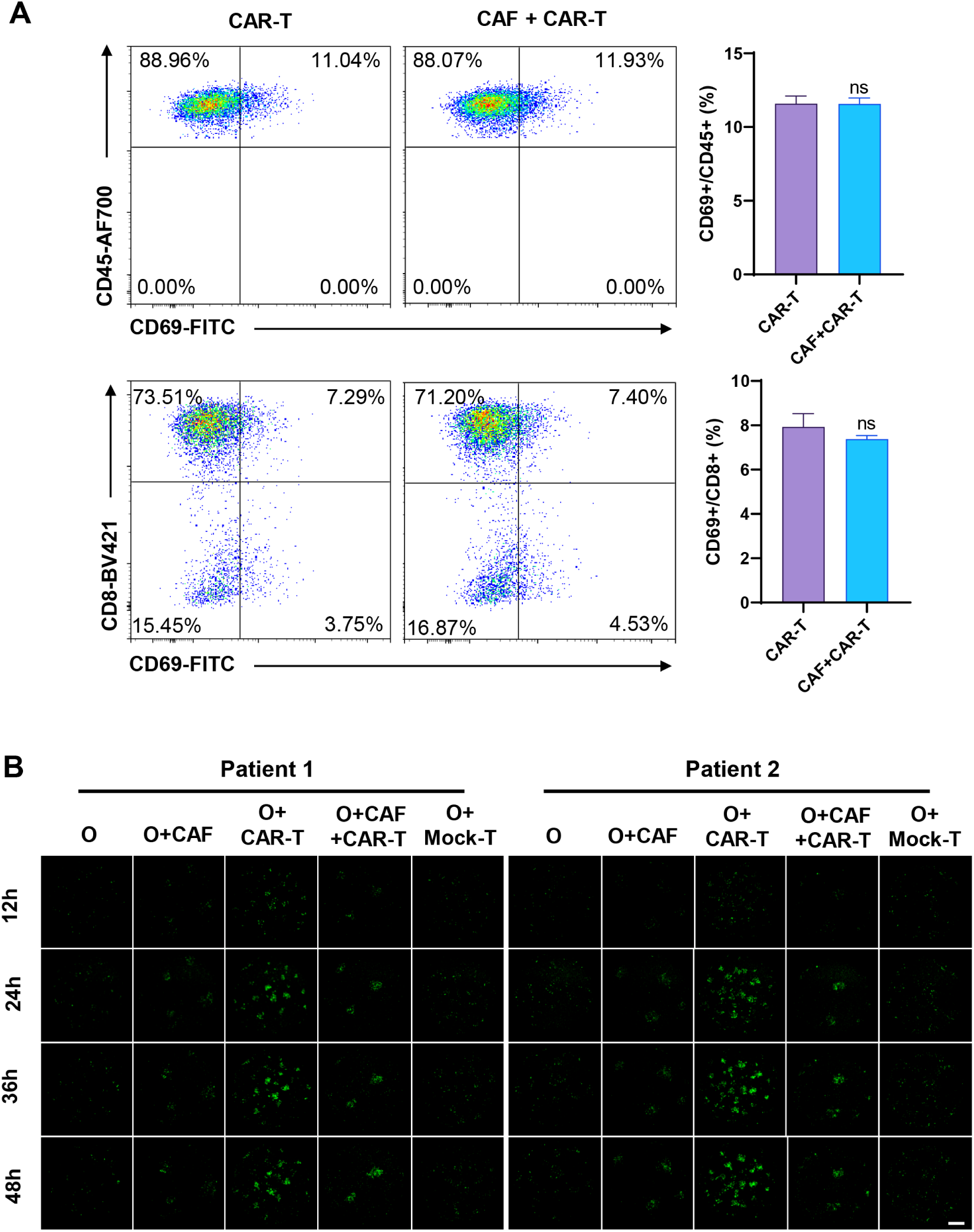
Mono-culture and CAF co-culture platform for CAR-T cell expansion and activation and antitumor efficacy assessment, Related to Figure 4. (A) Flow cytometry results and bar graphs illustrating the quantification of immune cell populations in various cultivation groups from patient 1. The graphs in the top row show cell ratio of CD45+CAR+ population. The graphs in the bottom row further shows cell ratio of CD8+CAR+ population. (B) Representative images of caspase-3 staining (green) in organoids co-culture with or without CAFs models after co-culture with or without CAR-T cells during 12-48 hours. Scale bars represent 100 μm. Data are presented as mean ± SD. Significant differences between groups are indicated by asterisks (* = p < 0.05, ** = p < 0.01, *** = p < 0.001).

**Figure S4.**
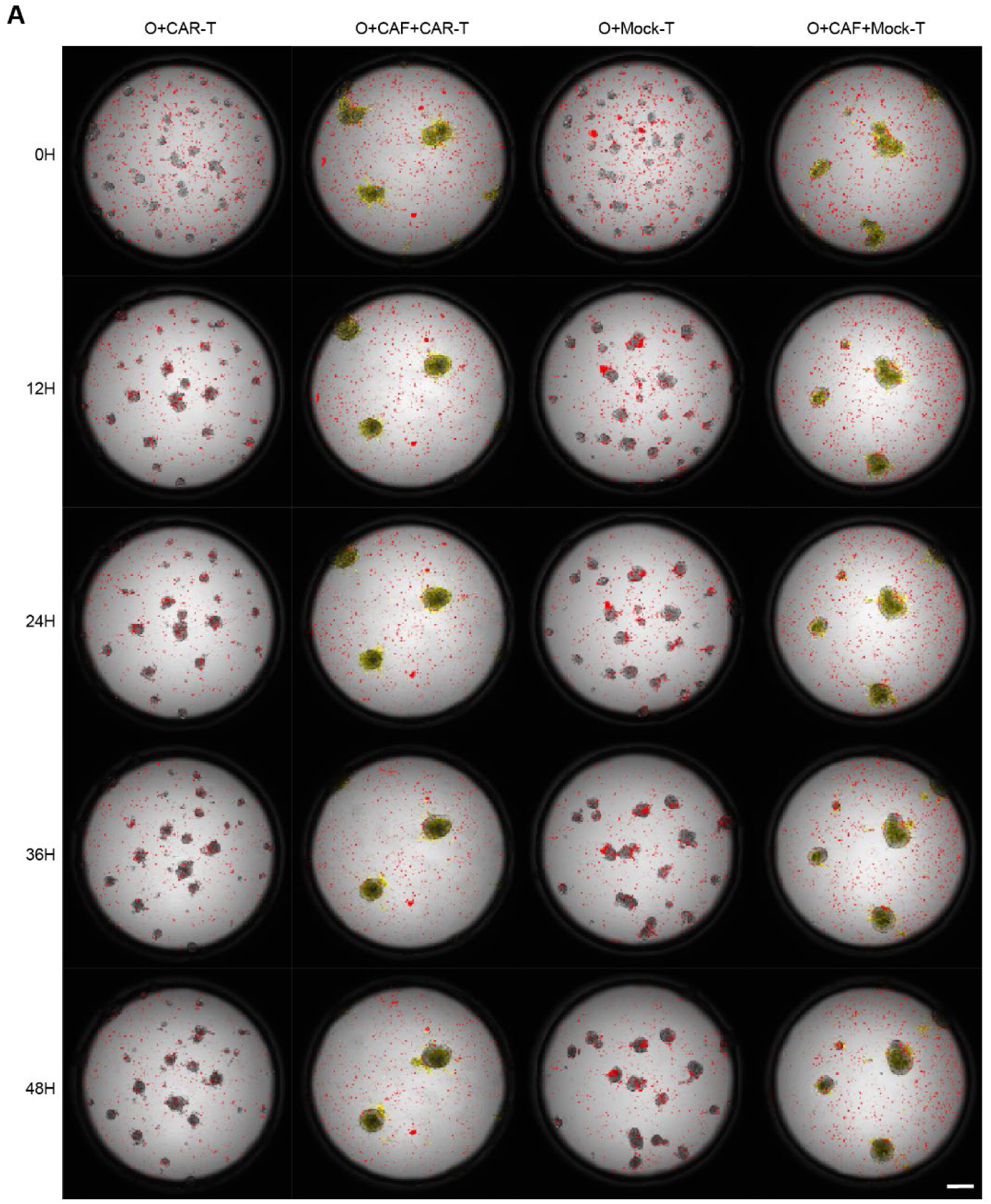
Time-lapse imaging of CAR-T cell infiltration into organoids, Related to Figure 5. CAR-T cells were added into mono-culture and co-culture models. CAR-T cells and CAF cells are respectively stained red and yellow. Organoids were visualized by phase contrast microscopy. Images show representative fields of view at 0, 12, 24, 36, and 48 hours post-co-culture. This demonstrates the effect of CAF cells on CAR-T cell infiltration and organoid interaction over time.

## Supplementary materialSupplementary video

Representative views of caspase-3 staining (green) in organoids after co-culture with CAR-T cells with or without CAFs during 6-24 hours.

